# Mosquitoes of the Maculipennis complex in Northern Italy

**DOI:** 10.1101/2020.11.02.365262

**Authors:** Mattia Calzolari, Rosanna Desiato, Alessandro Albieri, Veronica Bellavia, Michela Bertola, Paolo Bonilauri, Emanuele Callegari, Sabrina Canziani, Davide Lelli, Andrea Mosca, Paolo Mulatti, Simone Peletto, Silvia Ravagnan, Paolo Roberto, Deborah Torri, Marco Pombi, Marco Di Luca, Fabrizio Montarsi

## Abstract

The correct identification of mosquito vectors is often hampered by the presence of morphologically indiscernible sibling species. The Maculipennis complex is one of these groups which include both malaria vectors of primary importance and species of low/negligible epidemiological relevance, of which distribution data in Italy are outdated. Our study was aimed at providing an updated distribution of Maculipennis complex in Northern Italy through the sampling and morphological/molecular identification of specimens from five regions. The most abundant species was *Anopheles messeae s.l.* (2032), followed by *Anopheles maculipennis s.s.* (418), *Anopheles atroparvus* (28) and *Anopheles melanoon* (13). The distribution of species was characterized by Ecological Niche Models (ENMs), fed by recorded points of presence. ENMs provided clues on the ecological preferences of the detected species, with *An. messeae s.l.* linked to stable breeding sites and *An. maculipennis s.s.* more associated to ephemeral breeding sites. We demonstrate that historical *Anopheles* malaria vectors are still widespread in Northern Italy.

## Introduction

In early 1900, after the incrimination of *Anopheles* mosquito as a malaria vector, malariologists made big attempts to solve the puzzling phenomenon of “Anophelism without malaria”, that is, the absence of malaria in areas with an abundant presence of mosquitoes that seemed to transmit the disease in other areas (Kettle, 1995). The enigma was solved after the description of *Anopheles maculipennis sensu lato (s.l.)* as a species complex (i.e. the Maculipennis complex) which includes sibling species indistinguishable as imago, but differing in distribution, behaviour, ecology and in vectorial capacity for malaria (Kettle, 1995). Falleroni (1926) was the first to observe differences in the shape and exochorion patterns of eggs oviposited by engorged females. While partially overlapping between different species, for the first time this character allowed differentiation within the species complex. Attempts to find other diagnostic characters to separate the species complex were undertaken with particular attention to chaetotaxy in larvae and male genitalia. However, these characters frequently partially overlap between different species (Severini *et al.*, 2009), making morphological identification inconclusive. The molecular barcoding technique overcomes these morphological inconsistencies by sequencing evolutionary conserved loci. The internal transcribed spacer 2 (ITS2) marker proved particularly useful for discerning species in mosquito complexes (Bebe, 2018).

Two of the main malaria vectors in historical times in Italy, *Anopheles labranchiae* Falleroni, 1926, and *Anopheles sacharovi* Favre, 1903, belonged to the Maculipennis complex (Severini *et al*., 2009). In Italy, the complex also enumerated *Anopheles atroparvus* Vann. Thiel, 1927, which was considered to be a secondary vector in Italy, yet an important vector in northern Europe (Tekken & Knols, 2007). Other species present were *Anopheles maculipennis sensu stricto* (*s.s.*) Meigen, 1818, *Anopheles melanoon* Hakett, 1934, *Anopheles messeae* Falleroni, 1926, and the presumptive species *Anopheles subalpinus* Hackett & Lewis, 1935, now synonymised with *An. melanoon* (Linton *et al.*, 2002, Boccolini *et al.*, 2003). Other species in the complex were described in the Palearctic region based on cytogenetic (e.g. polytene chromosome inversions) (Artemov *et al.*, 2018; Andreeva *et al.*, 2007; Naumenko *et al.*, 2020) and barcoding approaches (Nicolescu *et al.*, 2004; Gordeev *et al.*, 2005; Djadid *et al.*, 2007): *Anopheles beklemishevi,* Stegnii & Kabanova, 1976, *Anopheles martinius,* Shingarev, 1926, *Anopheles artemievi,* Gordeyev, Zvantsov, Goryacheva, Shaikevich & Yezhov, 2005, *Anopheles persiensis,* Linton, Sedaghat & Harbach, 2003, *Anopheles daciae,* Linton, Nicolescu & Harbach, 2004, although the validity of some of these taxa is still debated. For instance, the species *An. messeae* was tentatively split into two species (*An. messeae* and *An. daciae)* (Nicolescu *et al.*, 2004) but the hierarchic status of these taxa within the complex is controversial. We overcame this critical issue by referring to the taxon *An. messeae* in a broader sense as *An. messeae s.l.*

Studies which started in the 1920s characterized the distribution of the Italian species (Bietolini *et al.*, 2006): *An. labranchiae* was present in central and southern Italy, while *An. atroparvus* was reported in northern regions. *Anopheles messeae* and *An. maculipennis s.s.* were recorded along the Italian peninsula, as was *An. melanoon* which, however, was characterized by a fragmentary presence. *Anopheles sacharovi* was reported in Italy but it seems to have disappeared following malaria eradication campaigns. Its last recorded date in Northern Italy dates back to 1959 (Romi *et al.*, 1997; Zamburlini & Cargnus, 1998).

Malaria was endemic in several areas of Italy until the 1950s, when the disease was eradicated following a strong campaign against mosquitoes and changes in people’s lifestyles (Gratz, 2006); the WHO finally declared the country free from malaria in the late 1970s (Zahar, 1990). Where mosquito vectors are present, locally-acquired malaria cases still occur after the arrival of affected persons, as reported in Greece since 2009 (NPHO, 2019). Moreover, areas with relevant residual malariogenic potential are still present in Italy (Romi *et al.*, 2012) and several cases of cryptic malaria were recorded in this country from 1997 until 2017 (Baldari *et al.*, 1998; Romi *et al.*, 2012; Boccolini *et al.*, 2020). In this scenario, the characterization of species distribution is strongly recommended to support a correct risk assessment (Boccolini *et al.*, 2020; ECDC, 2017).

After the eradication of malaria, the interest in anopheline fauna characterization decreased progressively and little data were available to corroborate the historical distribution of malaria vectors in northern Italy. The first aim of this study is to fill this knowledge gap by an extensive field sampling and identifying the Maculipennis complex species present by means of the barcoding technique. Furthermore, we used obtained presence points to model the environmental suitability of the surveyed territory for the different species detected, providing a useful tool for assessing the possible health risk associated with these vectors. The Ecological Niche Models (ENM) obtained provide interesting insights about characterizing different ecological traits of the species complex.

## Results

We collected more than 23,000 mosquitoes of the Maculipennis complex from the Po Valley, the widest plain in Italy. A large part of these mosquitoes was collected through attraction traps in fixed sites, which were sampled several times during the favourable season, i.e. between June and September. We used the geometric mean of Maculipennis complex mosquitoes collected per night of sampling in these sites in 2017-18 as a proxy for their abundance (Figure 1). We molecularly characterized 2,490 mosquitoes of the Maculipennis complex: 2,031 *An. messeae s.l.* (81.6%), 418 *An. maculipennis s.s.* (16.8%), 28 *An. atroparvus* (1.1%) and 13 *An. melanoon* (0.5%) (Figure 2, Table 1, Table S1). Of these mosquitoes, 17 specimens were collected as larvae in 5 sites, 846 were collected in 101 sites with manual aspirations specifically performed to collect them, and 1,627 were sampled through attraction traps in 198 sites, mainly within the frame of West Nile Virus (WNV) surveillance plans. The majority of tested mosquitoes were collected in 2017 and 2018, 1,046 and 1,277 respectively, while 125 specimens were collected before 2017 and 42 in 2019 (details in Table S2).

**Figure 1.**
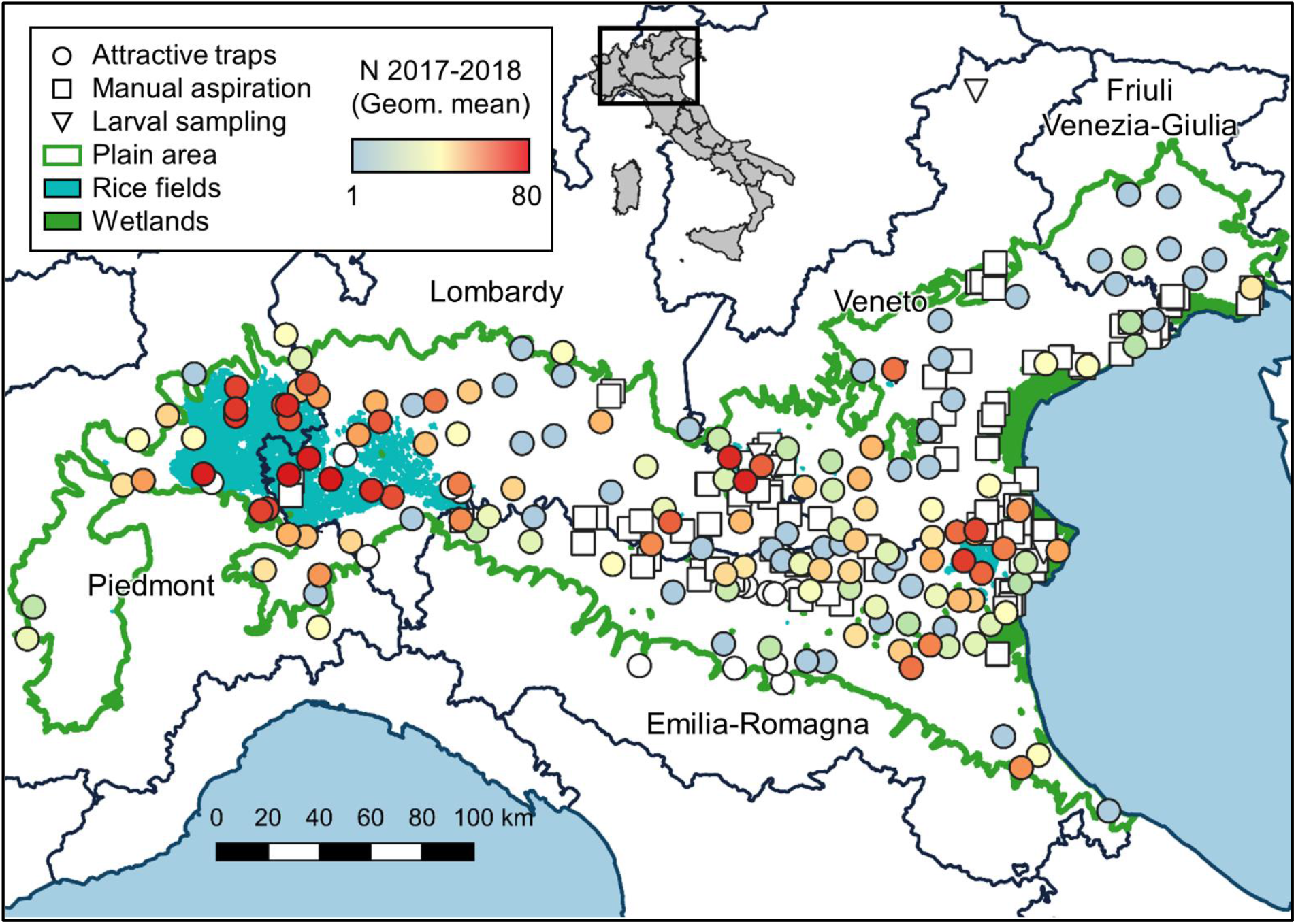
Location of sampled sites on the Northern Italy map, with reference to the boundaries of the Pianura Padana, type of sampling and the geometric mean of mosquitoes of the Maculipennis complex collected per night in the 2017-18 seasons.

**Figure 2.**
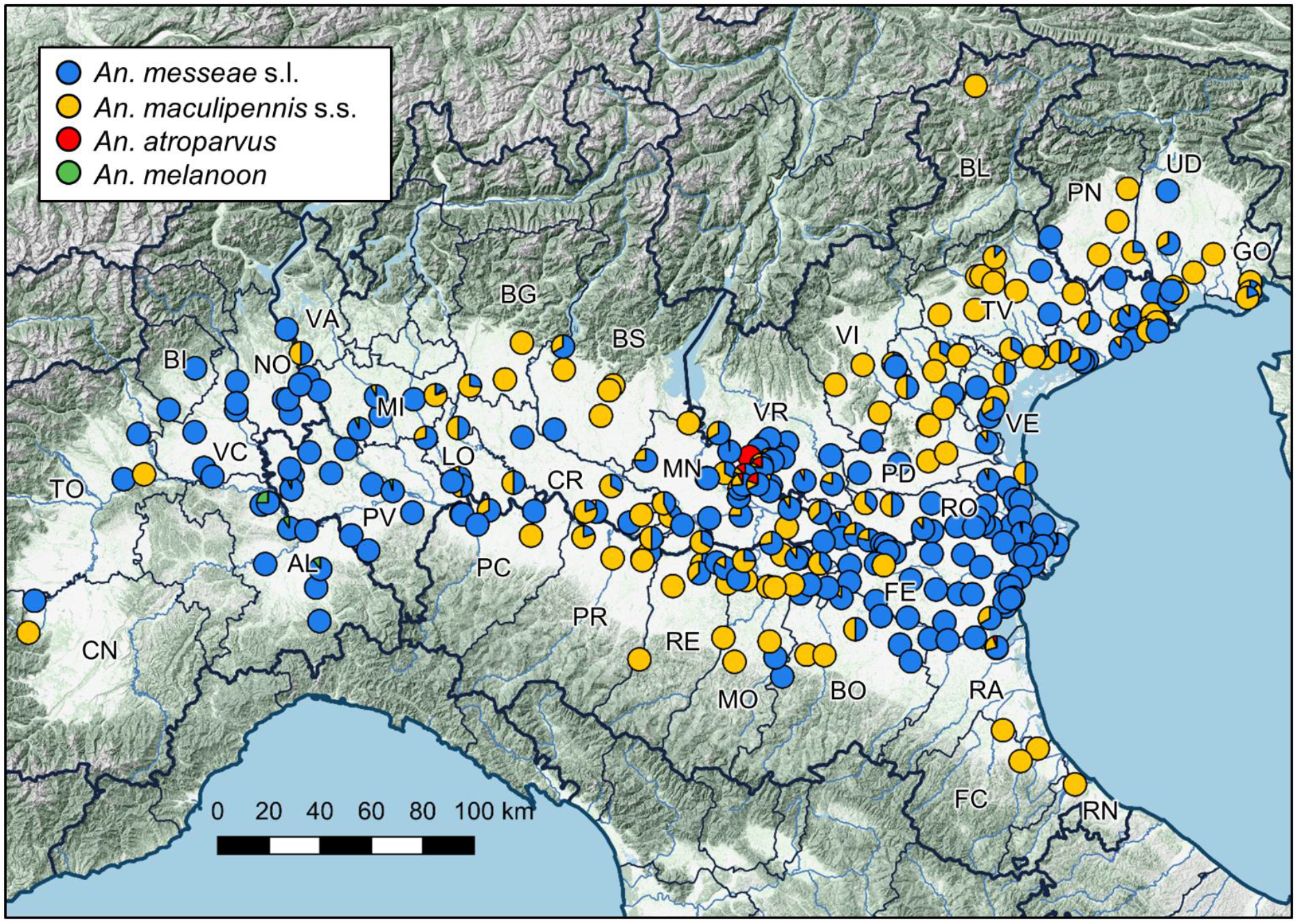
Proportion of different species identified in sampled sites. Abbreviations of Italian provinces (NUT3) with identified specimens are shown.

**Table 1.** Mosquitoes of the Maculipennis complex identified at species level with reference to the region of provenance and number of sites in which they were sampled.

|  | PIE |  | LOM |  | E-R |  | VEN |  | FVG |  | Total |  |
| --- | --- | --- | --- | --- | --- | --- | --- | --- | --- | --- | --- | --- |
|  | N | Sites | N | Sites | N | Sites | N | Sites | N | Sites | N | Sites |
| *An. messeae* s.l. | 281 | 24 | 617 | 50 | 408 | 68 | 709 | 84 | 16 | 7 | 2031 | 232 |
| *An. maculipennis* s.s. | 2 | 2 | 117 | 37 | 85 | 42 | 167 | 66 | 47 | 12 | 418 | 159 |
| *An. atroparvus* |  |  | 17 | 4 | 1 | 1 | 10 | 3 |  |  | 28 | 8 |
| *An. melanoon* | 6 | 3 | 7 | 6 |  |  |  |  |  |  | 13 | 9 |
| Total | 289 |  | 758 |  | 494 |  | 886 |  | 63 |  | 2490 |  |

PIE Piedmont, LOM Lombardy, E-R Emilia-Romagna, VEN Veneto, FVG Friuli Venezia-Giulia

### Barcoding

We obtained 2,330 ITS2 sequences (Table 2) which allowed for 93.6% of the achieved identification, confirming the discriminatory power of this marker for species complexes. The remaining 160 specimens were identified by Cytochrome C Oxidase-I (COI) (33) sequencing or by real-time PCR (127).

**Table 2.** ITS2 haplotypes recorded in this study for mosquito species, with reference to the number of specimens and accession number

| Species<br>(polymorphic bases) | Haplo | EBI AN | PIE | LOM | E-R | VEN | FVG |
| --- | --- | --- | --- | --- | --- | --- | --- |
| *An. messeae s.l.*<br>(161 165 167) |  |  |  |  |  |  |  |
|  | TTC | LR898482 | 161 | 392 | 239 | 79 | 2 |
|  | WTC | LR898483 | 63 | 102 | 84 | 202 | 5 |
|  | WTY | LR898484 | 15 | 77 | 38 | 258 | 3 |
|  | WWY | LR898485 | 21 | 32 | 4 | 44 |  |
|  | TTY | LR898486 |  | 6 | 24 | 5 |  |
|  | ATC | LR898487 |  | 1 | 2 |  |  |
|  | WTT | LR898488 |  | 1 |  |  |  |
|  | TWY | LR898490 |  | 1 |  |  |  |
|  | ATY | LR898491 |  | 1 |  |  |  |
|  | AWT | LR898482 |  | 1 |  |  |  |
| *An. maculipennis s.s.*<br>(one haplotype) |  |  |  |  |  |  |  |
|  |  | LR898492 | 2 | 115 | 81 | 130 | 38 |
| *An. atroparvus*<br>(136 138 147) |  |  |  |  |  |  |  |
|  | GGA | LR898493 |  | 3 |  |  |  |
|  | RRR | LR898494 |  | 7 |  | 1 |  |
|  | RRA | LR898495 |  | 4 | 1 | 9 |  |
|  | ARR | LR898496 |  | 2 |  |  |  |
|  | RGA | LR898497 |  | 1 |  |  |  |
| *An. melanoon*<br>(301) |  |  |  |  |  |  |  |
|  | T | LR898498 | 6 | 5 |  |  |  |
|  | W | LR898499 |  | 2 |  |  |  |
|  |  |  | 328 | 753 | 473 | 728 | 48 |

Haplo haplotype, AN Accession Number, PIE Piedmont, LOM Lombardy, E-R Emilia-Romagna,
VEN Veneto, FVG Friuli Venezia-Giulia

The ITS2 sequences obtained in this study were aligned with homologue sequences of Italian species of the complex available in GenBank and employed to obtain the phylogenetic tree in Figure 3. The sequences were grouped into four well-supported clades with conspecific reference sequences, enabling the unambiguous classification of our specimens in the four taxa: *An. maculipennis s.s., An. messeae s.l., An. melanoon, An. atroparvus.*

All ITS2 sequences were referred to a specific haplotype comprising sequences with ambiguous bases (Table 2), due to the presence of unequal double peaks systematically recorded in specific positions of electropherograms (Figure S1). We obtained a univocal haplotype, 292 bp long (excluding the 5.8 S and 28 S motifs in the obtained amplicon sequences) for the species *An. maculipennis s.s.,* identical to the available public sequences. We observed polymorphisms among sequences ascribable to *An. messeae s.l.* (305 bp long) *An. atroparvus* (307 bp) and *An. melanoon* (302 bp). We observed this variable pattern in three sites of *An. messeae s.l.* sequences, as previously reported (Lilja *et al.*, 2020; Naumenko *et al*., 2020), in three sites of *An. atroparvus* and one site in *An. melanoon* (Table 2). This latter site was the same site already reported as inter-individually polymorphic (Di Luca *et al.*, 2004). We did not observe other differences in obtained sequences falling into the same clade. This is also true for *An. messeae s.l.,* where all sequences showed the same pattern with A in position 362 and C in position 382, the two positions considered diagnostic for the identification of *An. daciae* (Lilja *et al.*, 2020; Naumenko *et al.*, 2020).

**Figure 3.**
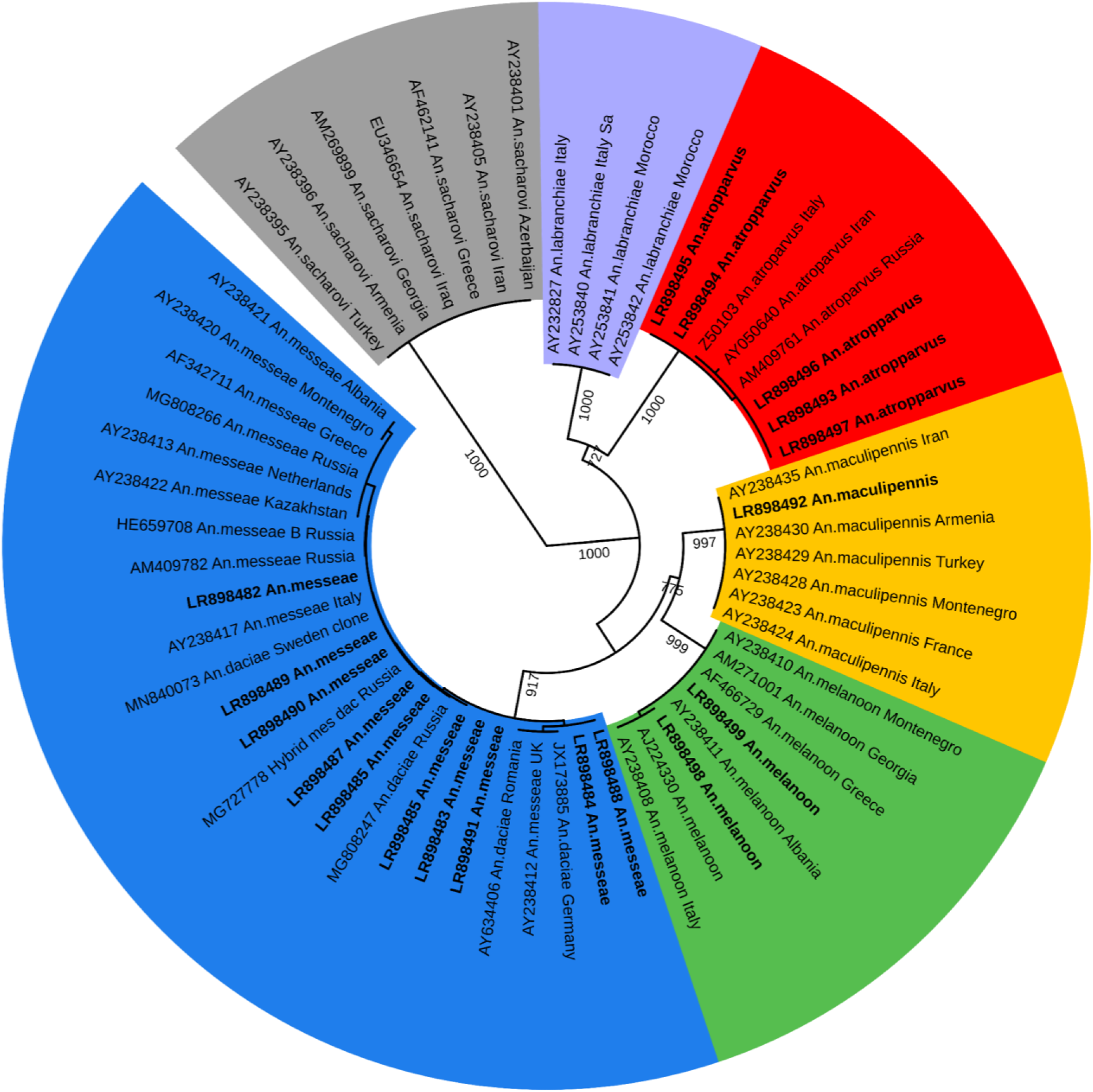
Maximum likelihood tree obtained by ITS2 haplotypes from this work (in bold with EBI accession numbers) and reference sequences from GenBank (accession number reported). Bootstrap values on 1000 replicates reported near the node, only values over 700 are shown.

### Ecological niche modelling

Following the screening of 148 available variables, a set of 34 were selected to be included in the Ecological Niche Models (ENM) (Table 3). As the number of capture sites for *An. atroparvus* and *An. melanoon* were scarce (eight and nine for the two species respectively) (Figure 4, top panel), ENMs were only built for *An. messeae s.l.* and *An. maculipennis s.s.* Overall performance for the ENMs was good, with AUC values of 0.80 for *An. messeae s.l.,* and 0.74 for *An. maculipennis s.s.*

**Table 3.**
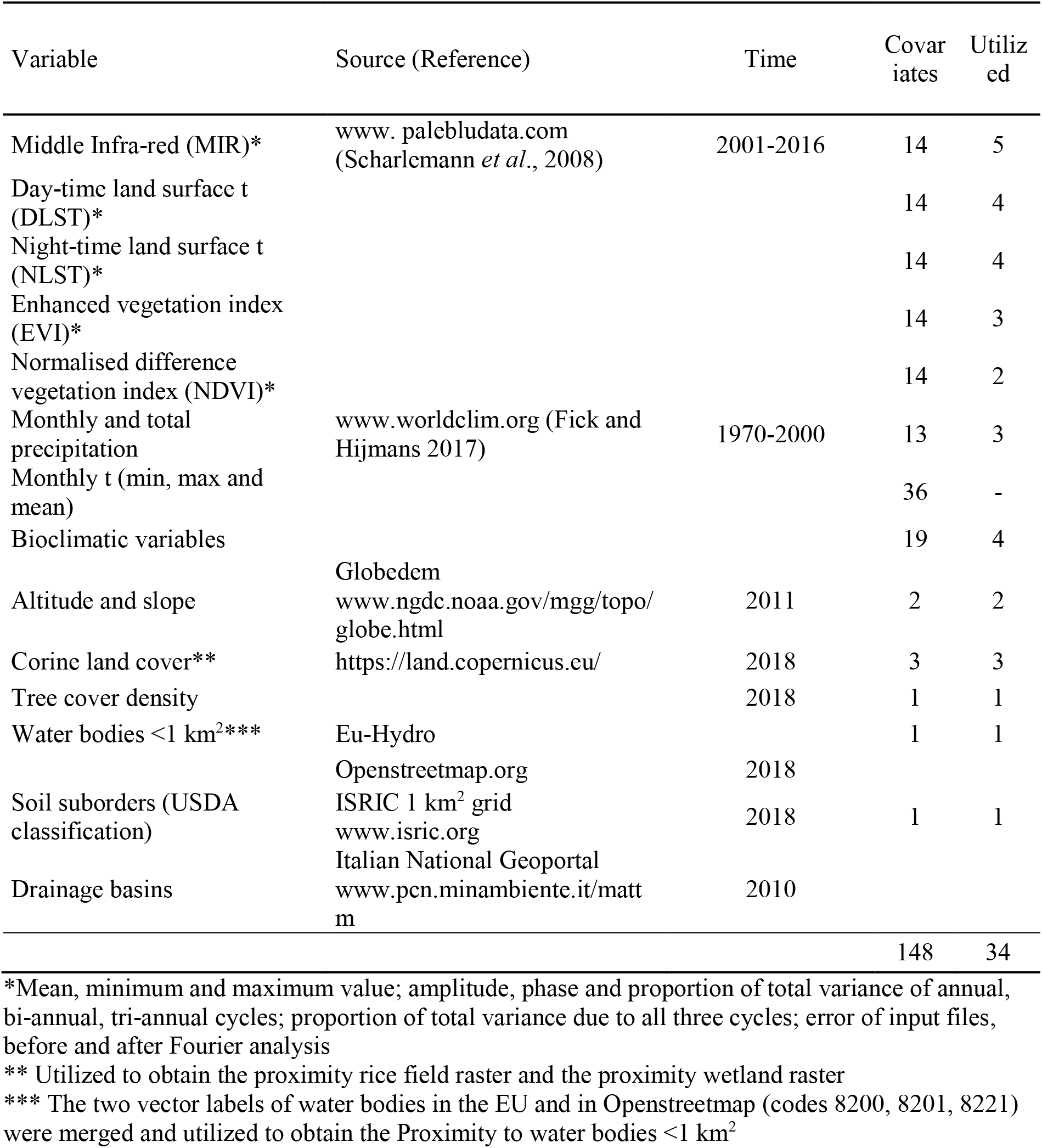
List of screened covariates with reference to those utilized for implementing the models.

| Variable | Source (Reference) | Time | Covariates | Utilized |
| --- | --- | --- | --- | --- |
| Middle Infra-red (MIR)* | www.palebludata.com<br>(Scharlemann *et al.*, 2008) | 2001-2016 | 14 | 5 |
| Day-time land surface t (DLST)* |  |  | 14 | 4 |
| Night-time land surface t (NLST)* |  |  | 14 | 4 |
| Enhanced vegetation index (EVI)* |  |  | 14 | 3 |
| Normalised difference vegetation index (NDVI)* |  |  | 14 | 2 |
| Monthly and total precipitation | www.worldclim.org (Fick and Hijmans 2017) | 1970-2000 | 13 | 3 |
| Monthly t (min, max and mean) |  |  | 36 | - |
| Bioclimatic variables |  |  | 19 | 4 |
| Altitude and slope | Globedem<br>www.ngdc.noaa.gov/mgg/topo/globe.html | 2011 | 2 | 2 |
| Corine land cover** | https://land.copernicus.eu/ | 2018 | 3 | 3 |
| Tree cover density |  | 2018 | 1 | 1 |
| Water bodies <1 km<sup>2</sup>*** | Eu-Hydro |  | 1 | 1 |
|  | Openstreetmap.org | 2018 |  |  |
| Soil suborders (USDA classification) | ISRIC 1 km<sup>2</sup> grid<br>www.isric.org | 2018 | 1 | 1 |
| Drainage basins | Italian National Geoportal<br>www.pcn.minambiente.it/mattm | 2010 |  |  |
|  |  |  | 148 | 34 |

\*Mean, minimum and maximum value; amplitude, phase and proportion of total variance of annual, bi-annual, tri-annual cycles; proportion of total variance due to all three cycles; error of input files, before and after Fourier analysis
\*\* Utilized to obtain the proximity rice field raster and the proximity wetland raster
\*\*\* The two vector labels of water bodies in the EU and in Openstreetmap (codes 8200, 8201, 8221) were merged and utilized to obtain the Proximity to water bodies <1 km<sup>2</sup>

The environmental suitability map suggested higher probabilities of the presence of *An. messeae s.l.* in the proximity of large breeding sites (Figure 4): westward near the rice fields in the Piedmont and Lombardy Regions, and close to wetlands. This is also confirmed by the relative contribution of the environmental variables to the ENM (Table 4), with distance to potential breeding sites *(Proximity to rice fields, Proximity to water bodies < 1 km^2^,* and *Proximity to wetlands)* providing a cumulative contribution of 41.4%. In addition, covariates related to elevation proved to be a relevant contribution to define the presence/absence of the species (Table 4, *Altitude* and *Slope;* cumulative contribution: 15.5%). *Proximity to rice fields* also resulted in being the covariate with the highest model gain when used alone; moreover, when the factor was omitted, the model had the lowest gain. This indicates that the covariate provided the most important pieces of information for defining the presence of *An. messeae s.l.*

**Table 4.** Relative contributions to the ENMs of selected covariates, with the related confidence intervals (CI).

| Covariate | *An. messeae* s.l. |  | *An. maculipennis* s.s. |  |
| --- | --- | --- | --- | --- |
|  | % contribution | CI | % contribution | CI |
| Proximity to rice fields | 25.7 | 27.0-24.5 | 1.7 | 1.8-1.6 |
| Altitude | 7.3 | 9.2-5.4 | 19.9 | 22.2-17.5 |
| Slope | 8.2 | 9.7-6.7 | 7.3 | 9.2-5.3 |
| Proximity to water bodies <1 km<sup>2</sup> | 8.8 | 9.4-8.2 | 5.9 | 6.9-4.9 |
| Proximity to wetlands | 6.9 | 8.3-5.5 | 5.7 | 6.7-4.7 |
| Soil type (USDA classification) | 2.6 | 2.9-2.3 | 9.5 | 12.1-6.9 |
| Middle infra-red (phase of tri-annual cycle) | 2.7 | 3.3-2.2 | 6.7 | 7.8-5.7 |
| Corine land cover | 7.0 | 7.6-6.4 | 2.1 | 2.5-1.7 |
| October precipitations | 2.6 | 3.3-2.0 | 6.1 | 6.8-5.3 |
| NDVI (minimum value) | 2.2 | 2.8-1.6 | 6.2 | 7.1-5.3 |

The most suitable area for *An. maculipennis s.s.* resulted located in the central and northern part of the Po Valley (Figure 4). For this species, the contribution of proximity to breeding sites was less important (13.3%), while covariates related to altitude and slope resulted more relevant (27.2% contribution in total) (Table 3). However, soil type also showed a marked contribution in itself (9.5%), with a higher relevance of moisty soils characterized by the likely presence of surface water and a negative influence of arid soils on the model (Figure S2). Also, factors describing the effect of vegetation (NDVI and middle-red reflectance) appeared to have an important cumulative contribution (12.9% in total) in the ENM for *An. maculipennis,* in marked contrast to what was observed for *An. messeae s.l.* (cumulative contribution 4.9%), indicating different vegetation requirements for the two species. The covariate with the highest model gain when used by itself for *An. maculipennis s.s.* was *altitude*, indicating that altitude alone could provide the most useful information when defining the presence of the species. On the contrary, the greater loss of gain in a multi-variable model was observed if *Proximity to water bodies <1 km^2^* was omitted which, therefore, appeared to be the factor bearing the most important information that is not captured by the other covariates included in the ENM.

The contribution to the models of all employed covariates is reported in Table S3.

**Figure 4.**
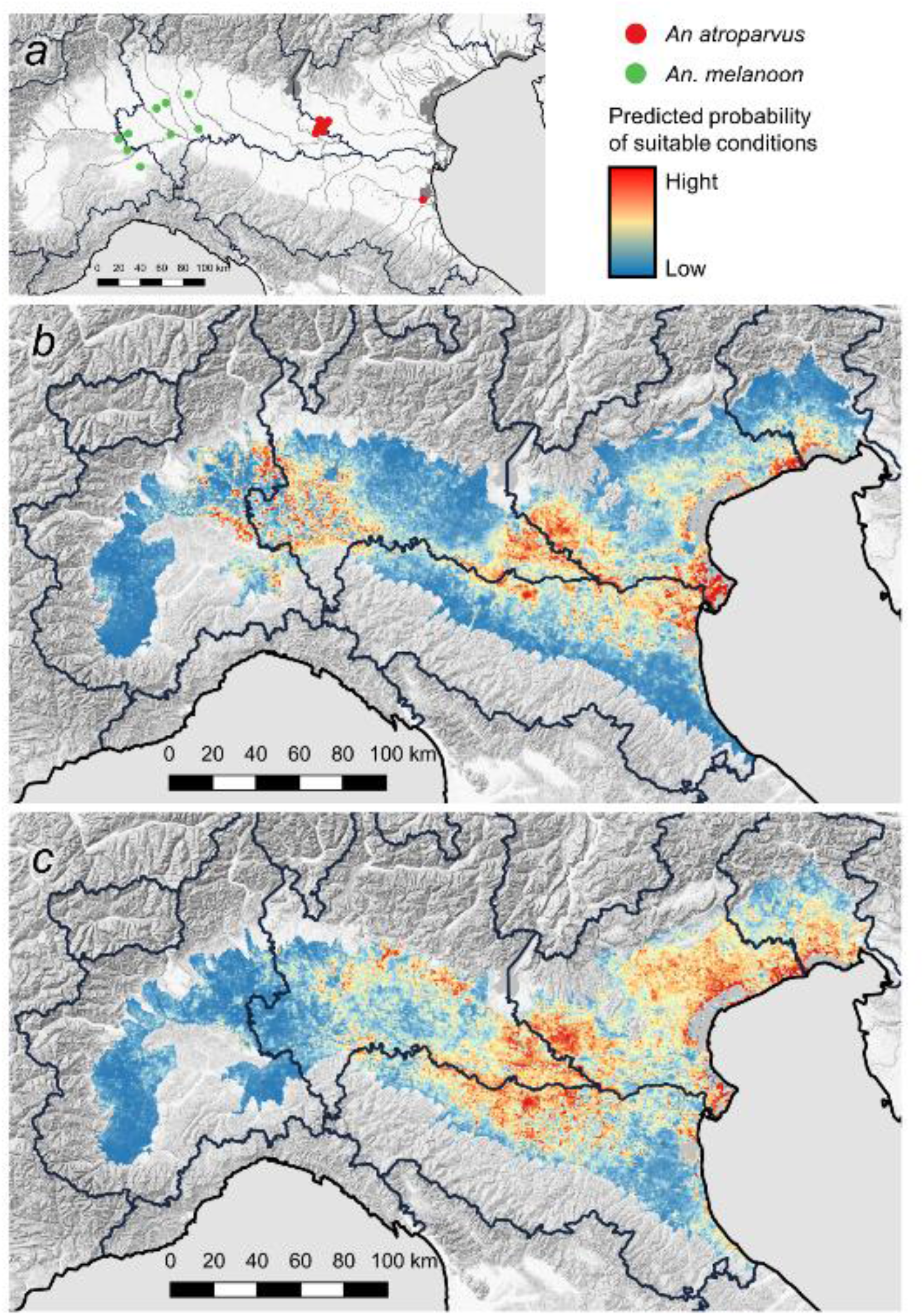
a) Site of detection of *An. atroparvus* (red) and *An. melanoon* (green). Output of ENM obtained by Maxent for *An. messeae s.l.* (b) and *An. maculipennis s.s.* (c).

## Discussion

This study provides detailed data on the distribution of species of the Maculipennis complex in Northern Italy, filling the gap in the knowledge gathered over the last decades.

The use of the ITS2 marker allowed the identification of *An. messeae s.l., An. maculipennis s.s.* and the rarer *An. melanoon* and *An. atroparvus,* suggesting the disappearance of *An. sacharovi* in the surveyed area.

While ITS2 sequences ascribable to *An. maculipennis s.s.* were all identical, we recorded an unexpected variability in some positions of the ITS2 sequences in other species. The ITS2 marker is widely used for low intraspecific variability, but it could be present in hundreds of copies in a single genome (Bebe, 2018), and these copies can differ at individual level for single mutations, insertions or deletions, as extensively reported in mosquitoes (Batovska *et al.*, 2017). The presence of a single mutation in the same specimen systematically caused a double peak of different intensity in a particular position of the electropherogram. We recorded these multiple peaks in three positions of *An. messeae s.l.,* three positions of *An. atroparvus* and one position of *An. melanoon*. These variable bases, caused by an intra-individual polymorphism, are not reliable as diagnostic markers for definition at species level. While these particular polymorphisms were already reported in *An. messeae s.l.,* we reported these polymorphic sites in another two species of the Maculipennis complex, *An. melanoon* and *An. atroparvus*.

The ITS2 polymorphic sites were used to define new species. The taxon *An. daciae* was earlier defined through the presence of five polymorphisms in *An. messeae* from Romania (Nicolescu *et al.*, 2004). As observed in our data and demonstrated by cloning (Lilja *et al.*, 2020), three of these polymorphisms were intra-individual, making only the last two sites reliable for biomolecular identification. In the last two polymorphic sites, our specimens show the same haplotype, strongly suggesting that we faced a single taxonomic entity (species or form) in the study area. Assuming the validity of *An. daciae*, we did not detect *An. messeae* in Northern Italy, but only *An. daciae*.

The taxonomy of *An. messeae s.l.* is puzzling: the existence of two taxa, named A and B chromosomal forms, was already suggested (Novikov & Kabanova, 1979). The identity of form A with *An. daciae* was then postulated (Vaulin & Novikov, 2012), but the rank of the two taxa, *An. daciae/An. messeae* or A/B forms, is still debated. The described intragenomic heterogeneity raised doubts on the use of ITS2 as a univocal species marker for the definition of *An. daciae* (Bezzhonova & Goryacheva, 2008). Besides, specimens bearing both variants in the two above-mentioned diagnostic sites were reported (Bezzhonova & Goryacheva, 2008; Vaulin & Novikov, 2012; Naumenko *et al.*, 2020), strongly suggesting the same extent of interbreeding between *An. daciae* and *An. messeae* (Naumenko *et al*., 2020), and unequivocal diagnostic morphological traits were not identified between the two taxa (Nicolescu *et al.*, 2004). On the other hand, significantly different relative proportions of chromosomal inversions were observed between A and B forms (Novikov & Shevchenko, 2001). Also, a fixed chromosomal inversion (X1) was observed in *An. messeae*, in addition to a relevant genetic distance between *An. messeae* and *An. daciae*, mainly on the X chromosome (Naumenko *et al.*, 2020). We suggest that a critical review of the *An. messeae* status, with more robust evidences based on strong genetic and ecological data, is needed before unambiguously splitting *An. messeae* into two species.

Moreover, the definition of the taxa of the Maculipennis complex is a difficult task, as highlighted by the absence of clear morphological differences and the incomplete reproductive barriers among members of the complex (Kitzmiller *et al.*, 1967). This is probably due to the recent, or incipient, process of speciation which pose several taxa of the complex in the “grey zone” (De Queiroz, 2007) that are difficult to characterize. But the definition of taxonomic relationship inside the complex is a stimulating task, because *Anopheles* complexes are an ideal context to define the boundaries of sibling species (Mallet *et al.*, 2016; Pombi *et al.*, 2017; Fouet *et al.*, 2017; Alquezar *et al.*, 2018).

The elucidation of relations inside this complex is not only a taxonomic curiosity, since the identification of a new species can have strong epidemiological implications. Ecological (as suitable breeding sites) and behavioural (as host preferences) characteristics of different species may affect the capacity to transmit pathogens, such as *Plasmodium* parasites (Manguin *et al.*, 2008; Cohuet *et al.*, 2010). The “anophelism without malaria” phenomenon demonstrated the epidemiological importance of finely characterizing species in the complex. Part of the specimens used were obtained by exploiting existing surveillance plans, mainly targeting theWNV, also demonstrating the usefulness of these plans for tasks that are outside of the main scope. Although traps used in WNV surveillance targeting *Culex* mosquitoes were not expressly studied for trapping *Anopheles* mosquitoes, they collected a relevant number of the latter. Moreover, we regularly monitored WNV sites for two years, fortnightly over the summer season, thus obtaining comparable data on the abundance of *Anopheles* from monitored sites. This data demonstrated that mosquitoes of the Maculipennis complex are more abundant in areas rich in large breeding sites (such as rice fields and wetlands).

Even if a comparison with historical data is not easy, due to the different sampling strategies and identification protocols, our data suggest that *An. melanoon* and *An. atroparvus* are less widespread than in the past. Only few specimens of *An. atroparvus*, a historical malaria vector in northern Europe, were collected in the eastern part of the area, supporting evidence of the scarce presence of the species that seems rarer than in the past, particularly in the Po Delta area. The same scenario is described for *An. melanoon*, recorded in the western part of the monitored area, which seems to have reduced its diffusion when compared to the past (Bietolini *et al*., 2006).

The presence of the more abundant species *An. messeae s.l.* and *An. maculipennis s.s.* appeared to be more likely in plain areas, as demonstrated by the importance of altitude and slope in defining environmental suitability. Nevertheless, the two species showed a different geographical distribution. While *An. messeae s.l.* resulted in having a higher chance of occurrence in western and eastern parts of the monitored area where large breeding sites were also reported, *An. maculipennis s.s.* appeared more diffused in the central section of the study area where larger breeding sites are rare and many of these are transient (Figure 2). In fact, the two species seem to show different preferences in breeding site typologies. *Anopheles messeae s.l.* was more related to consistent breeding sites such as rice fields, wetlands and sparse water bodies. *An. maculipennis s.s.* seems to exploit more ephemeral breeding sites, as indicated by the relevance in the ENM of slope and soil types characterized by the propensity to host surface waters and then linked to the presence of transient breeding habitats. These results, obtained by modelling, are consistent with the two species’ known preferences, with *An. messeae* preferring large water bodies with irregular water levels rich in vegetation, and *An. maculipennis s.l.* occupying small water bodies with scarce vegetation and a widely-fluctuating daily temperature (Becker *et al.*, 2010).

The presence of these potential malaria vectors highlights epidemiological importance for an understanding of their distribution, a fundamental prerequisite to evaluate the risk of malaria introduction via Plasmodia carriers. The distribution of potential vectors differed from the past and is constantly changing according to ecological mutations, such as land use modifications and climatic change. The presence of competent vectors, together with favourable conditions, can lead to the occurrence of locally-acquired malaria, as demonstrated in Italy in 1997 (Baldari *et al.*, 1998), and more recently and impressively in Greece with 109 cases classified as locally-acquired from 2009 to 2019 (NPHO, 2019). A deadly malaria case that involved a child in Italy in 2017, and which successively proved to be nosocomial, aroused a great deal of public opinion (Boccolini *et al.*, 2020; ECDC, 2017). Previous knowledge of the *Anopheles* fauna would have provided the basic data for a first rapid assessment of the case’s origin and a prompt and clear risk communication to the people. Species of the Maculipennis complex exhibit a different vector competence for Plasmodia. Furthermore, Plasmodium strains that can be potentially imported are completely different to those that historically circulated in the country and are no longer present. Experimental infections and xenodiagnoses demonstrated the ability of European *An. messeae, An. atroparvus*, and *An. sacharovi* to transmit Asian *Plasmodium vivax* (Daskova & Rasnicyn, 1982). While European *An. atroparvus,* and *An. labranchiae* seem refractory to African strains of *Plasmodium falciparum* (de Zulueta *et al.*, 1975; Ramsdale & Coluzzi, 1975; Teodorescu *et al.*, 1978; Daskova & Rasnicyn, 1982; Sousa, 2008; Toty *et al.*, 2010; Romi *et al.*, 2012; van Dorp *et al.*, 2020).

In order to assess the risk of locally-acquired malaria cases, these fragmentary data must be corroborated by comprehensive studies to characterize the vectorial competence of local mosquitoes, particularly for new taxa, when they have been described with exotic and potentially importable strains of Plasmodia. All these data, although difficult to obtain, are necessary to improve our preparedness to face emergencies.

## Materials and Methods

### Study area

The surveyed area included the widest plain in Italy, the Po Valley (Pianura Padana or Pianura Padano-Veneta), which includes parts of the Piedmont, Lombardy, Emilia-Romagna, Veneto and Friuli-Venezia Giulia regions. This area, of about 46,000 km^2^, enumerates more than 20 million inhabitants and is the most densely populated in Italy, with some of the largest Italian cities (Milan, Turin, Bologna, and Venice). This territory is geared to agriculture, characterized by intensive farming and animal husbandry, with few hedges, rare scattered trees and a dense irrigation network. Industrial settlements and residential areas frequently interweave this agricultural environment. Natural areas are rare, mainly represented by river borders, characterized by riparian vegetation, or protected and re-naturalized areas.

The same section of the surveyed area presents widespread presence of breeding sites suitable for *Anopheles* mosquitoes, such as the rice field areas (2,250 km^2^ between eastern Piedmont and western Lombardy, 110 km^2^ between the Ferrara and Rovigo provinces and 40 km^2^ between the Mantua and Verona provinces, besides other smaller areas). These areas were endemic for malaria in the past, particularly in the eastern part bordering the Adriatic Sea, at the Po River Delta. Potential mosquito breeding sites such as restored natural areas, quarries, irrigation channels and natural and artificial wetlands are abundant and sparsely present in the Po Valley.

### Entomological collection

Mosquito samples belonging to the Maculipennis complex were collected mainly in 2017 and 2018 by active sampling or attraction traps, in addition to stored samples from previous years (Table S2). A large part of tested mosquitoes was retrieved from entomological monitoring in the frame of the WNV surveillance, in which adult mosquitoes were sampled by traps (CDC-like) baited with dry ice as a source of carbon dioxide in fixed stations of the Po Valley (Calzolari *et al.*, 2015). Mosquitoes derived from entomological samplings performed for other purposes were identified to a lesser extent (e.g. *Leishmania infantum* surveillance, nuisance monitoring), sometimes sampled by CDC-light traps (15 specimens). Sites surveyed for WNV were sampled fortnightly from June to September for two seasons. The abundance of specimens belonging to the Maculipennis complex was expressed as the geometric mean of mosquitoes in the complex per sampling (Figure 1). To complete the picture, we actively collected larvae and adult *Anopheles* mosquitoes in areas where WNV traps were unsuccessful. The survey was carried out by actively collecting larvae by dipping and adult mosquitoes by direct aspiration in resting sites, from May to October. These sites included animal shelters (bovines, equines, goats and poultry) in farms to collect engorged and host-seeking mosquitoes, and uninhabited buildings to collect mosquitoes ready for overwinter.

We separated Maculipennis complex specimens from other mosquitoes according to morphological keys (Severini *et al.*, 2009; Becker *et al.*, 2003). Sites and relative methods of sampling are depicted in Figure 1.

### Barcoding and sequence analysis

We selected part of the collected specimens and analysed a maximum of 13 specimens belonging to the Maculipennis complex per sample (namely a collection made in one site in one day). We used the entire bodies or just one leg of these mosquitoes for the biomolecular analysis; in this case, the rest of the body was individually stored in a cryotube at −80 °C. We extracted the DNA which was submitted to traditional PCR protocol for amplification of the internal transcribed sequences 2 (ITS2) according to Marinucci *et al.*, 1999; this PCR amplified the DNA encoding for ITS2 and part of ribosomal 5.8 S and 28 S rRNA subunits. The obtained amplicons were then sequenced. Part of the samples were identified by sequencing of the Cytochrome C Oxidase-I (COI) marker (Jalali *et al.*, 2015) or real-time PCR (Lühken *et al.*, 2016).

We defined all obtained haplotypes, also comprising ambiguous bases, and ascribed to them all the sequences obtained in a Region (Table 2). In order to prevent the incorporation of possible sequencing errors, we set the arbitrary threshold of 1% of conspecific sequences to insert ambiguous bases in one haplotype. These sequences were used in a BLAST search to obtain a homologue sequence of vouchered specimens from different countries. Only the ITS2 part of obtained sequences was aligned with the MAFFT algorithm (Katho *et al.*, 2019). We used the alignment to infer a phylogenetic tree by means of the maximum likelihood method implemented in PhyML 3.1 software (Guindon & Gascuel, 2019) with 1,000 bootstrap replicates. The model TPM2+G was selected according to the jModelTest2 software (Darriba *et al.*, 2012). The tree was visualized with ITol software (Letunic & Bork, 2019). These representative sequences were deposited in the European Nucleotide Archive (EBI) database under accession numbers from LR898482 to LR898499;.

### Ecological niche modelling

We used the presence points of the detected species to model the environmental suitability of the different species in the surveyed area according to the maximum entropy approach, using MaxEnt software v3.4.1 (Phillips *et al.*, 2017). This software estimates environmental suitability for a species departing from a set of occurrence locations and gridded covariates, maximising the entropy in geographic space or, in other terms, minimizing relative entropy between covariates (Elith *et al.*, 2011). A series of meteorological (such as temperatures and precipitations) and ecological variables (such as soil type and indexes) or covariates, were obtained from different sources (Table 3). Proximity maps were obtained by utilizing vector files for different categories of breeding sites (rice fields, wetlands, sparse water bodies under 1 km^2^ of surface and riverbeds).

All covariates were acquired as raster, then rescaled and aligned at 1 km^2^ spatial resolution in ASCII format on the surveyed area extent (WGS84 projection) by using Qgis v3.10. We defined the reference raster area by enlarging the boundaries of the plain area by 15 km, represented as the vector file described in Calzolari *et al.*, (2015).

An explorative analysis was run with all covariates. Those with no contribution in the models, and thus ecologically irrelevant, were excluded from the final analysis. We screened covariates for correlation, selecting those having a Pearson’s correlation coefficient of less than 0.7. In order to avoid potential collinearity problems within the set of covariates used in the model, we assessed collinearity with the variance inflation factor (VIF) which is a measure of correlation between pairs of covariates. Correlation and collinearity analysis were performed in R version 3.5 and ENM Tools (Warren *et al.*, 2010). The set of covariates used in the models is reported in the supplemental material (Table S3).

Ten replicates were done with the cross-validation run and the cloglog model output grid format. In order to overcome the different sampling efforts in different areas, a bias file was utilized categorizing provinces into three groups (100%, 60% and 40%) according to densities of observations.

Modelling performance was evaluated using the Area Under Curve (AUC) value. AUC measures the model’s sensitivity and was used to test the model’s performance with real observations in the training area. An AUC value of 0.5 shows that the model predicts randomly, while a value close to 1 indicates optimal model performance. AUC > 0.75 was considered as a good value (Philips & Dudik, 2008).

The better covariates’ relative contributions to the MaxEnt model were estimated using percent contribution. Covariate contribution was tested by Jackknife analysis in the MaxEnt model to get alternate estimates of the most important covariates in the model (Philips *et al.*, 2006). We performed a GIS analysis with QGIS v3.10 software.

## Acknowledgements

Authors would thank all field and laboratory technicians who collaborated to this study.

## Supplementary material

**Table S1.**
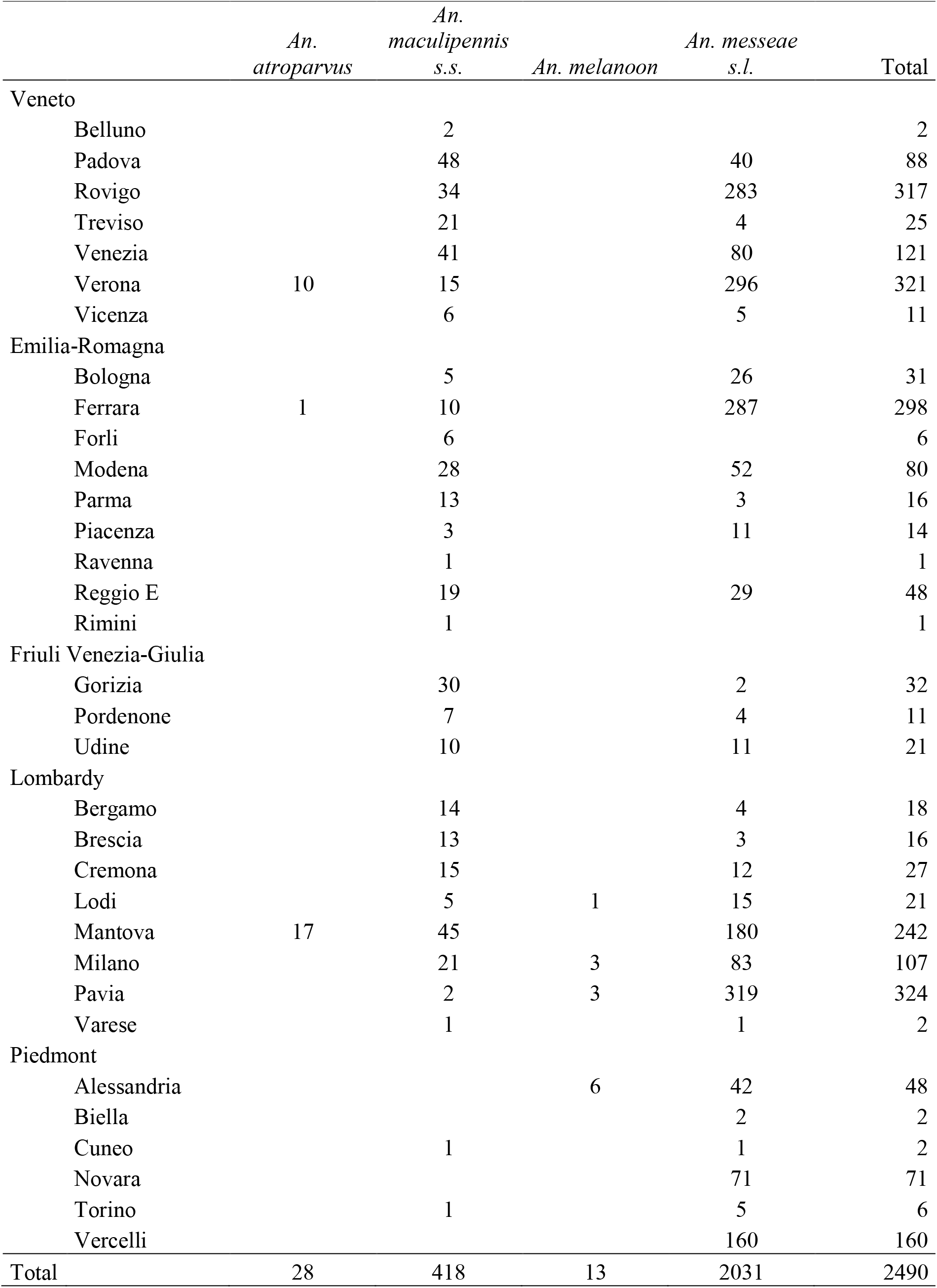
Mosquitoes of the Maculipennis complex identified at provincial level (NUT3).

|  | *An.*<br>*atroparvus* | *An.*<br>*maculipennis*<br>*s.s.* | *An. melanoon* | *An. messeae*<br>*s.l.* | Total |
| --- | --- | --- | --- | --- | --- |
| Veneto |  |  |  |  |  |
| Belluno |  | 2 |  |  | 2 |
| Padova |  | 48 |  | 40 | 88 |
| Rovigo |  | 34 |  | 283 | 317 |
| Treviso |  | 21 |  | 4 | 25 |
| Venezia |  | 41 |  | 80 | 121 |
| Verona | 10 | 15 |  | 296 | 321 |
| Vicenza |  | 6 |  | 5 | 11 |
| Emilia-Romagna |  |  |  |  |  |
| Bologna |  | 5 |  | 26 | 31 |
| Ferrara | 1 | 10 |  | 287 | 298 |
| Forli |  | 6 |  |  | 6 |
| Modena |  | 28 |  | 52 | 80 |
| Parma |  | 13 |  | 3 | 16 |
| Piacenza |  | 3 |  | 11 | 14 |
| Ravenna |  | 1 |  |  | 1 |
| Reggio E |  | 19 |  | 29 | 48 |
| Rimini |  | 1 |  |  | 1 |
| Friuli Venezia-Giulia |  |  |  |  |  |
| Gorizia |  | 30 |  | 2 | 32 |
| Pordenone |  | 7 |  | 4 | 11 |
| Udine |  | 10 |  | 11 | 21 |
| Lombardy |  |  |  |  |  |
| Bergamo |  | 14 |  | 4 | 18 |
| Brescia |  | 13 |  | 3 | 16 |
| Cremona |  | 15 |  | 12 | 27 |
| Lodi |  | 5 | 1 | 15 | 21 |
| Mantova | 17 | 45 |  | 180 | 242 |
| Milano |  | 21 | 3 | 83 | 107 |
| Pavia |  | 2 | 3 | 319 | 324 |
| Varese |  | 1 |  | 1 | 2 |
| Piedmont |  |  |  |  |  |
| Alessandria |  |  | 6 | 42 | 48 |
| Biella |  |  |  | 2 | 2 |
| Cuneo |  | 1 |  | 1 | 2 |
| Novara |  |  |  | 71 | 71 |
| Torino |  | 1 |  | 5 | 6 |
| Vercelli |  |  |  | 160 | 160 |
| Total | 28 | 418 | 13 | 2031 | 2490 |

**Table S2.** Mosquito of the Maculipennis complex with reference to the year and method of sampling.

| Year | Sampling method | *An. atroparvus* | *An. maculipennis s.s.* | *An. melanoon* | *An. messeae s.l.* | Total |
| --- | --- | --- | --- | --- | --- | --- |
| 2011 | Trap |  | 5 |  | 2 | 7 |
| 2013 | Trap |  | 10 |  | 38 | 48 |
| 2014 | Trap |  | 5 |  | 6 | 11 |
| 2015 | Trap |  | 14 |  | 19 | 33 |
| 2016 | Trap |  |  |  | 26 | 26 |
| 2017 | Trap | 8 | 103 | 4 | 618 | 733 |
|  | Aspiration | 10 | 45 |  | 258 | 313 |
| 2018 | Trap |  | 103 | 7 | 617 | 727 |
|  | Aspiration | 10 | 128 | 1 | 394 | 533 |
|  | Larval sampling |  | 2 |  | 15 | 17 |
| 2019 | Trap |  | 3 |  | 39 | 42 |
| Total |  | 28 | 418 | 13 | 2031 | 2490 |

**Table S3.** Selected covariates after collinearity screening and their averaged contribution (Av.) to the obtained models and related confidence interval (CI).

| Species | Code | *An. messeae s.l.* |  | *An. maculipennis s.s.* |  |
| --- | --- | --- | --- | --- | --- |
|  |  | Av. | CI | Av. | CI |
| Precipitation of Wettest Quarter | bio16 | 0.4 | 0.6-0.2 | 0.3 | 0.5-0.1 |
| Isothermality* | bio3 | 0.4 | 0.7-0.1 | 0.6 | 1.0-0.3 |
| Min Temperature of Coldest Month | bio6 | 0.7 | 1.0-0.5 | 0.1 | 0.1-0.0 |
| Temperature Annual Range | bio7 | 0.4 | 0.6-0.2 | 1 | 1.2-0.8 |
| Corine Land Cover | clc2018 | 7 | 7.6-6.4 | 2.1 | 2.5-1.7 |
| Middle Infra-red, amplitude of annual cycle | er1603a1 | 2.5 | 3.0-2.1 | 1.3 | 1.6-0.9 |
| Middle Infra-red amplitude of tri-annual cycle | er1603a3 | 0.5 | 0.6-0.3 | 0.5 | 0.7-0.3 |
| Middle Infra-red, proportion of total variance due to annual cycle | er1603d1 | 1.1 | 1.2-0.9 | 1.7 | 2.2-1.3 |
| Middle Infra-red, proportion of total variance due to bi-annual cycle | er1603d2 | 1 | 1.1-0.8 | 3 | 3.5-2.5 |
| Middle Infra-red, phase of tri-annual cycle | er1603p3 | 2.7 | 3.3-2.2 | 6.7 | 7.8-5.7 |
| Day-time land surface temperature, amplitude of tri-annual cycle | er1607a3 | 1.4 | 2.1-0.8 | 1.2 | 1.6-0.8 |
| Day-time land surface temperature, proportion of total variance due to tri-annual cycle | er1607d3 | 0.3 | 0.6-0.1 | 0.9 | 1.2-0.5 |
| Day-time land surface temperature, % zero or above DN range | er1607e1 | 0.7 | 0.9-0.6 | 2.3 | 2.8-1.8 |
| Day-time land surface temperature, total variance | er1607vr | 1.5 | 2.0-1.0 | 0.1 | 0.2-0.1 |
| Night-time land surface temperature, amplitude of bi-annual cycle | er1608a2 | 1.6 | 2.1-1.0 | 1 | 1.4-0.7 |
| Night-time land surface temperature, amplitude of tri-annual cycle | er1608a3 | 1.3 | 1.8-0.8 | 1.1 | 1.4-0.9 |
| Night-time land surface temperature, minimum value | er1608mn | 0.6 | 0.8-0.3 | 0.2 | 0.4-0.1 |
| Night-time land surface temperature, total variance | er1608vr | 0.3 | 0.4-0.1 | 0.5 | 0.9-0.2 |
| Enhanced vegetation index (EVI), fourier mean for entire time series | er1614a0 | 1.5 | 1.8-1.1 | 2.5 | 3.4-1.7 |
| Enhanced vegetation index (EVI) Proportion of total variance due to annual cycle | er1614d1 | 1.4 | 1.7-1.1 | 1.9 | 2.4-1.4 |
| Enhanced vegetation index (EVI), total variance | er1614vr | 3.7 | 4.3-3.2 | 1 | 1.8-0.2 |
| Normalised difference vegetation index(NDVI) Amplitude of tri-annual cycle | er1615a3 | 1 | 1.2-0.8 | 1.4 | 1.7-1.0 |
| Normalised difference vegetation index(NDVI), minimum value | er1615mn | 2.2 | 2.8-1.6 | 6.2 | 7.1-5.3 |
| Digital elevation model (altitude) | globedem | 7.3 | 9.2-5.4 | 19.9 | 22.2-17.5 |
| Digital elevation model (slope) | slope | 8.2 | 9.7-6.7 | 7.3 | 9.2-5.3 |
| Proximity raster of small inland bodies | inland_os<br>m_proximity | 8.8 | 9.4-8.2 | 5.9 | 6.9-4.9 |
| Proximity raster of wetland | prox_clc4 | 6.9 | 8.3-5.5 | 5.7 | 6.7-4.7 |
| Proximity raster of river | proxy_rivers | 1.7 | 2.2-1.3 | 2.8 | 3.3-2.2 |
| October precipitations | rain10 | 2.6 | 3.3-2.0 | 6.1 | 6.8-5.3 |
| March precipitations | rain3 | 0.9 | 1.3-0.6 | 0.8 | 1.1-0.5 |
| July precipitations | rain7 | 0.4 | 0.6-0.3 | 1.4 | 1.7-1.1 |
| Proximity raster of ricefield | prox_ricefield | 25.7 | 27.0-24.5 | 1.7 | 1.8-1.6 |
| USDA soil classification | USDA<br>Soil | 2.6 | 2.9-2.3 | 9.5 | 12.1-6.9 |
| Tree cover | tree_cp_2015 | 0.6 | 0.7-0.5 | 1.2 | 1.7-0.7 |

**Figure S1.**
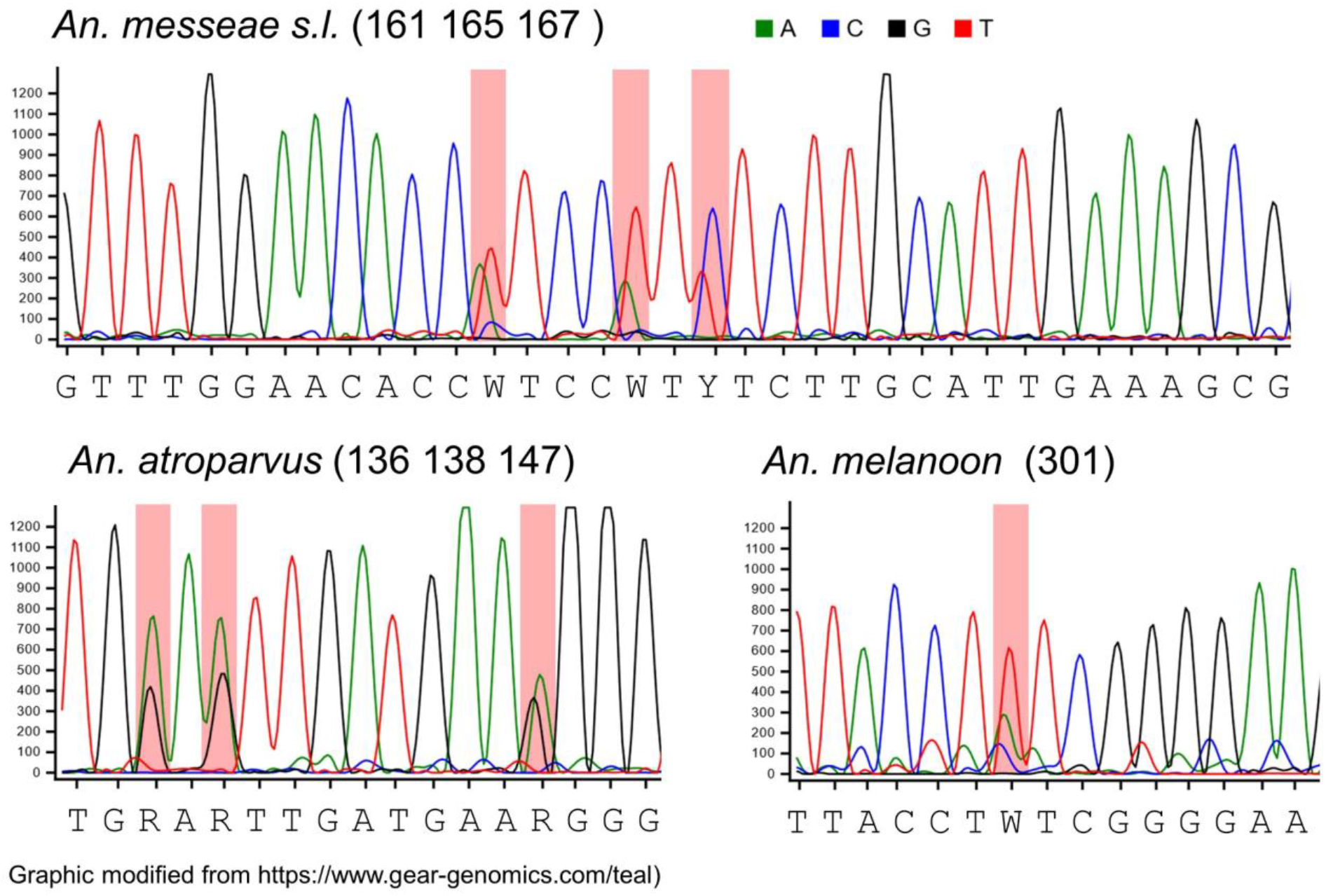
Examples of multiple peaks recorded in electropherograms.

**Figure S2.**
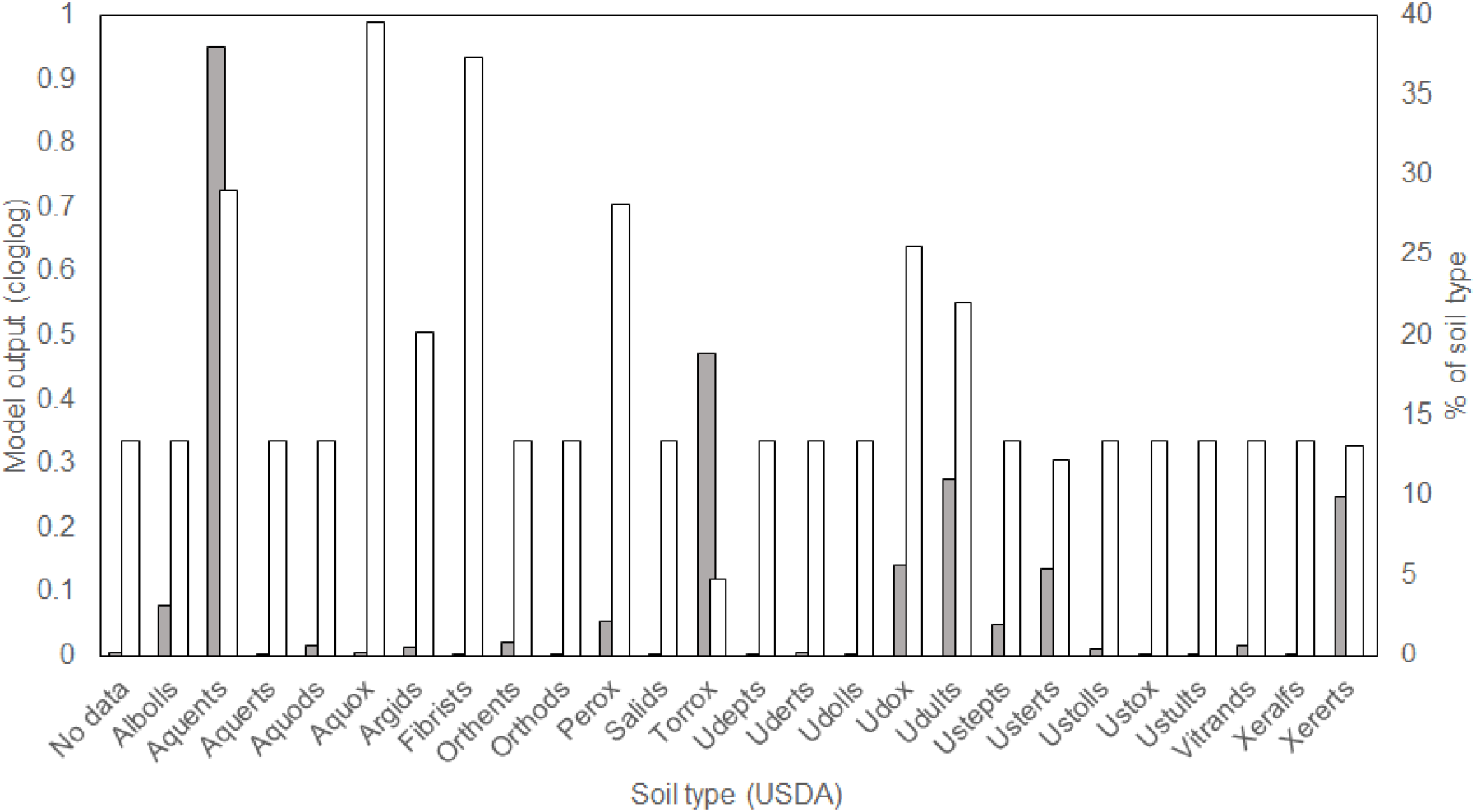
Relevance of soil classification in the ENM of *An. maculipennis s.s.* (white) with reference to the % area of different soil types in the surveyed area (grey).

